# Vascular and renal telomere shortening in mice exposed to chronic intermittent hypoxia

**DOI:** 10.1101/2021.02.25.432912

**Authors:** Mohammad Badran, Bisher Abuyassin, Najib Ayas, Don D. Sin, Ismail Laher

## Abstract

Obstructive sleep apnea (OSA) is a chronic condition characterized by chronic intermittent hypoxia (IH) and is associated with cardiovascular (CVD) and chronic kidney diseases (CKD). There is increased biomarkers of aging, such as telomere shortening, in patients with OSA. We assessed telomere lengths in aortic and renal tissues from mice exposed to 8 weeks of IH using a PCR protocol, and demonstrate significant telomere shortening in both tissues. This data indicates that IH, a hallmark of OSA, can accelerate vascular and renal aging that may contribute to OSA-induced CVD and CKD

## Introduction

Obstructive sleep apnea (OSA) is a chronic condition characterized by repetitive episodes of airway collapse inducing intermittent hypoxia (IH) and sleep fragmentation, and is considered an independent risk factor for many conditions such as cardiovascular (CVD) and chronic kidney diseases (CKD) (1,2). Prominent underlying pathological mechanisms involved in OSA-induced CVD and CKD are inflammation and oxidative stress (3). Telomere shortening, a biomarker of aging and cellular senescence that is influenced by oxidative stress (4), is associated with CVD and CKD (5,6). While telomere attrition in leukocytes has been documented in OSA patients (7), it is unknown whether the obstructive events leading to intermittent hypoxia can shorten telomeres in peripheral organs such as the aortic vasculature and kidneys. In this experiment, we exposed mice to 8 weeks of chronic intermittent hypoxia (IH) and measured telomere length in aortic and kidney tissues.

## Methods

### IH protocol

Male C57BL/6 mice (8-10 weeks old, 25-30 grams) were housed in customized cages placed in a room with a 12 h light-dark cycle (07:00 h-19:00 h) at a constant temperature (26°C). A gas control delivery system regulated the flow of room air and N_2_ into the customized cages to control the inspired O_2_ fraction (FiO_2_) levels in each cage. The level of FiO_2_ was reduced from 20.9% to 5.0% over a 30-s period and rapidly returned to room air levels using a burst of 100% O_2_ during the following 30-s period for 12hrs daily during the light cycle. This IH paradigm generates oxyhemoglobin nadir values in the 50-60% range. Mice were exposed to room air throughout the 12-h dark cycle (when mice are active). For the intermittent air (IA, control) group, normoxic gas was flushed periodically into the chambers over 24hrs at the same frequency as the protocol for inducing IH. Mice were exposed to either the chronic IH or intermittent air (control) for 8 weeks (8).

### Telomere length

The telomere lengths of mouse aortic and kidney tissues were measured using a qPCR protocol as described previously by Callicott and Womack (9). In brief, the assay measures the amount of telomeric DNA per sample as a ratio to the total amount of genomic DNA, which is given by the measurement of acidic ribosomal phosphorprotein PO (36B4) gene. Telomeric forward and reverse primer sequences (Sigma, The Woodlands, TX) were (5⍰→3⍰) CGG TTT GTT TGG GTT TGG GTT TGG GTT TGG GTT TGG GTT and (5⍰→3⍰) GGC TGG CCT TAC CCT TAC CCT TAC CCT TAC CCT TAC CCT, respectively, while murine 36B4 forward and reverse primer sequences were (5⍰→3⍰) ACT GGT CTA GGA CCC GAG AAG and (5⍰→3⍰) TCA ATG GTG CCT CTG GAG ATT, respectively. Measurements were performed in triplicates per sample on 384-well Clear Optical Reaction Plates (Applied Biosystems, Foster City, CA). A sample of DNA extracted from the lung tissue of a C57BL/6J mouse was added to each plate to serve as a reference. Calculations for telomere length over single copy gene ratio (T/S ratio) for each sample were compared to the T/S ratio of the reference DNA and expressed as relative telomere length.

## Results and Discussion

The effects of 8 weeks of exposure to IH on the telomere lengths in the aorta and kidneys of mice is shown in Fig.1. Exposing mice to 8 weeks of IH significantly shortened telomere lengths in aortic tissues (IH: 0.85 ± 0.02 kb/genome vs. IA: 0.92 ± 0.02 kb/genome, *p* = 0.028). Similarly, IH significantly shortened telomeres in renal tissues (IA: 0.77 ± 0.03 kb/genome vs. IH: 0.73 ± 0.01 kb/genome, *p* = 0.031).

This study provides novel data on the effects of IH, a hallmark of OSA, in accelerating the aging of aortic and kidney tissues as measured by telomere shortening. Our data provide a potential mechanism for the increased risk of CVD and CKD in patients with OSA. More mechanistic studies are needed to better evaluate the role of telomere shortening on other vascular tissues and to determine the potential benefits of treatments strategies such as continuous positive airway pressure (CPAP) in humans. It will also be important to design interventional studies to assess whether preserving telomere lengths can reduce the risk of CVD and CKD in patients with OSA by attenuating cellular replicative senescence in these key organs.

This study was funded by CIHR (Sleep Team Grant) and BC Lung Association

No financial disclosures were reported by the authors of this letter

---

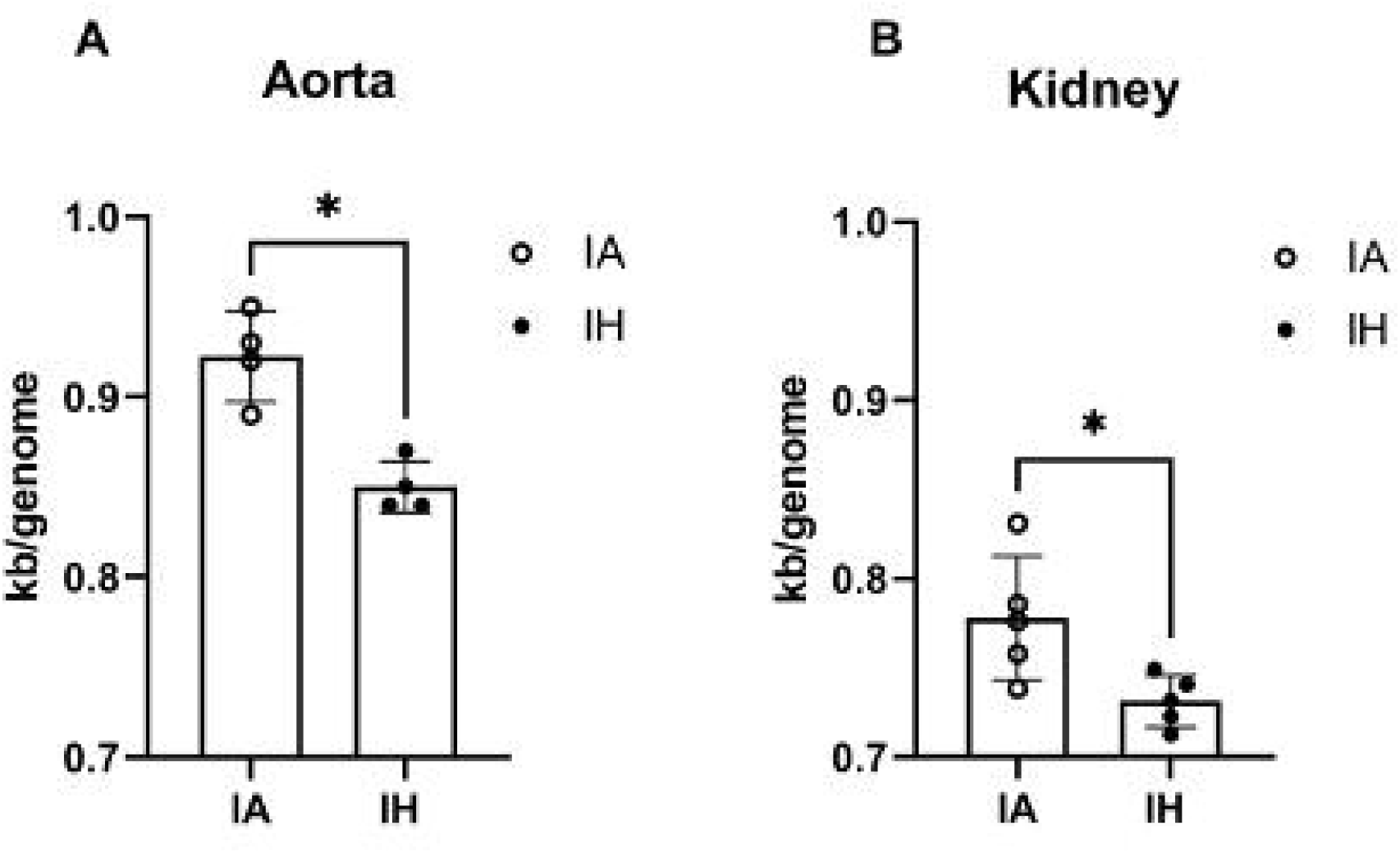

